# Assessing the reliability of an automated method for measuring dominance hierarchy in nonhuman primates

**DOI:** 10.1101/2020.11.23.389908

**Authors:** Sébastien Ballesta, Baptiste Sadoughi, Fabia Miss, Jamie Whitehouse, Géraud Aguenounon, Hélène Meunier

## Abstract

Among animals’ societies, dominance is an important social factor that influences inter-individual relationships. However, assessing dominance hierarchy can be a time-consuming activity which is potentially impeded by environmental factors, difficulties in the recognition of animals, or through the disturbance of animals during data collection. Here we took advantage of novel devices, Machines for Automated Learning and Testing (MALT), designed primarily to study nonhuman primates’ cognition - to additionally measure the social structure of a primate group. When working on a MALT, an animal can be replaced by another; which could reflect an asymmetric dominance relationship (or could happen by chance). To assess the reliability of our automated method, we analysed a sample of the automated conflicts with video scoring and found that 75% of these replacements include genuine forms of social displacements. We thus first designed a data filtering procedure to exclude events that should not be taken into account when automatically assessing social hierarchies in monkeys. Then, we analysed months of daily use of MALT by 25 semi-free ranging Tonkean macaques (*Macaca tonkeana*) and found that dominance relationships inferred from these interactions strongly correlate with the ones derived from observations of spontaneous agonistic interactions collected during the same time period. We demonstrate that this method can be used to assess the evolution of individual social status, as well as group-wide hierarchical stability longitudinally with minimal research labour. Further, it facilitates a continuous assessment of dominance hierarchies, even during unpredictable environmental or challenging social events. Altogether, this study supports the use of MALT as a reliable tool to automatically and dynamically assess social status within groups of nonhuman primates, including juveniles.

## Introduction

To build stable relationships, social animals, including primates, must respond appropriately to various social situations. This stability is indeed a significant aspect of social structure (Hinde 1976), and allows animals to prevent conflicts and to optimize their social relationships with others, both of which play a crucial role in individuals’ fitness (Silk 2007; Silk et al. 2010; Kulik et al. 2012; Majolo et al. 2012; McFarland and Majolo 2013; Kerhoas et al. 2014). Dominance is an important social factor that influences the daily interactions between group members in primates’ societies (Rowell 1974; Bernstein 1981). However, as the number of individuals in a social group increases, the number of interactions will also increase exponentially, which can make direct observations of social behaviors challenging and results in sparse data on dyadic relationships (de Vries 1995). Experimental methods involving a competitive context have been used in order to assess dominance hierarchy in non-human primates, however for optimal results, it may imply the use of water or food deprivation or require a behavioural training of the subjects (Hamilton 1960; Boelkins 1967; Christopher 1972; Clark and Dillon 1973; Wrangham 1981; Canteloup et al. 2016). In NHP (non-human primates), access to enrichments can also be considered to assess the social structure of the group (Chamove 1983; Ballesta et al. 2014). Although these methods are more time-efficient, these still require considerable human and time resources and may depend on the experimental context of competition (Brennan and Anderson 1988).

The fields of cognitive ethology and neuroscience have seen a recent increase in the development and use NHP, of Machines for Automated Learning and Testing (MALT) allowing the study of mental and social processes (Fagot and Bonté 2010; Gazes et al. 2013; Claidière et al. 2017; Fizet et al. 2017; Gazes et al. 2019; Gelardi et al. 2019). These modern protocols do not require isolating the subject from its social group and cognitive testing can be voluntarily performed, at their own pace, which improves animal welfare during data collection. These devices are a valuable refinement of the practices in cognitive ethology and may represent a change of paradigm in neuroscience that involve NHPs. Importantly, the behaviors and cognition of NHPs assessed by these devices are comparable to the one expressed in a laboratory setting (Gazes et al. 2013), and therefore extend computer-based accurate study of cognition to semi-free ranging animals. It is worth noting that these testing devices also represent a valuable environmental enrichment and contribute to increase the welfare of captive or semi-free ranging NHP (Bennett et al. 2016, 2018). So far, MALT have been identified for serving as a functioning tool in cognition research, while their potential to explore social dynamics in groups of NHPs provided with MALT has only started to be investigated (Claidière et al. 2017; Gelardi et al. 2019). Dominance in groups of NHP has so far been mostly studied using direct observation methods described by Altmann (Altmann 1974). These standard methods demonstrated their suitability for providing unbiased behavioral data and allowed gathering the vast majority of information we currently have on NHP sociality. However, in spite of their undeniable usefulness, these methods are time consuming, costly in terms of human resources, and limited regarding the quantity of data we can collect in a day. To overcome these limitations and explore a new potential of MALTs, one possibility is to use such automated devices to investigate social relationships and thus group structure. To evaluate the reliability of this method, we compare social information gathered through standard observation techniques with social information collected on the same social group automatically through MALTs. We analyzed 103,655 working sessions made by 25 Tonkean macaques (*Macaca tonkeana*) on four MALTs present at the Primate Center of the University of Strasbourg (Fizet et al. 2017). We observed that macaques can compete for access to the MALT by displacing other animals currently working on it. We therefore hypothesized that the outcome of these competitive interactions could inform us about the dominance hierarchy of the group, which was measured in parallel through direct observations in the macaques’ usual environment. In addition, as a proof of concept, we applied this method to depict the dynamic of the dominance hierarchy of the study group during a three-year period. We assessed the consequences of males removal on group stability and highlighted the usefulness of our method for group management of primates in captivity.

## Materials and Methods

### Subjects

We collected data on one social group of Tonkean macaques (*Macaca tonkeana*), all captive-born and housed at the Primate Center of the University of Strasbourg, France. Animals lived in semi-free ranging conditions in a wooded park of 3788m^2^ with permanent access to an indoor-outdoor shelter (2.5×7.5m - 2×4m). The group included 28 individuals with even sex ratio among adults (see **Table 1**), which is comparable to the composition of wild groups (Riley 2005, 2007). Individuals of less than 3 years old were considered as juvenile. Monkeys were fed with commercial primate pellets twice a day inside the indoor shelter and received fresh fruit and vegetables once a week outside observation hours. Water was provided *ad libitum* in the indoor shelter. Four females had contraceptive implants according to the Primate Center breeding program, and one female gave birth in February 2018. Out of the 28 individuals from the group, we collected data at the MALT from 25 individuals and data from direct observations on 23 individuals. The alpha male (determined by direct observations) of the group, ‘*Uly*’, never significantly engaged with the MALT for the last 4 years and therefore could not be included in our automated data collection. More data are needed in order to know if it is a personal specificity or a consequence of being the dominant individual in Tonkean macaques society (as this has not been observed in other species of monkeys (Claidière et al. 2017; Gazes et al. 2019; Gelardi et al. 2019). The other individual that never used the MALT was born in February 2018 (‘*Fic*’). The subject was considered too young to have an RFID chip in her forearms. We did not record a sufficient number of events (no data left after the filtering procedure described below) for the subject ‘*Wal*’ that was thus excluded from this analysis (see **Fig S1**). During direct observations, five subjects (‘*Bar*’, ‘*Ber*’, ‘*Ces*’, ‘Dor’ and ‘*Eri*’) were too young to be reliably identified in direct observations of social conflicts but were using the MALT at that time. Hence, the comparison of the dominance hierarchy obtained by automatic and observational data includes 22 out of the 28 individuals. Note that only after January 2019 ‘Bar’ and ‘Ber’ were old enough to be reliably identified during direct observations. Two key events could represent a disruption in the stability of the hierarchy: the 26th of May 2018, one adult male, (‘*Wot*’) and the 18th of January 2019, four adult males (‘*Yan*’, ‘*Yak*’, ‘*Wal*’, ‘*Wat*’) were removed for group-management purposes (Wooddell et al. 2017).

**Table 1.**
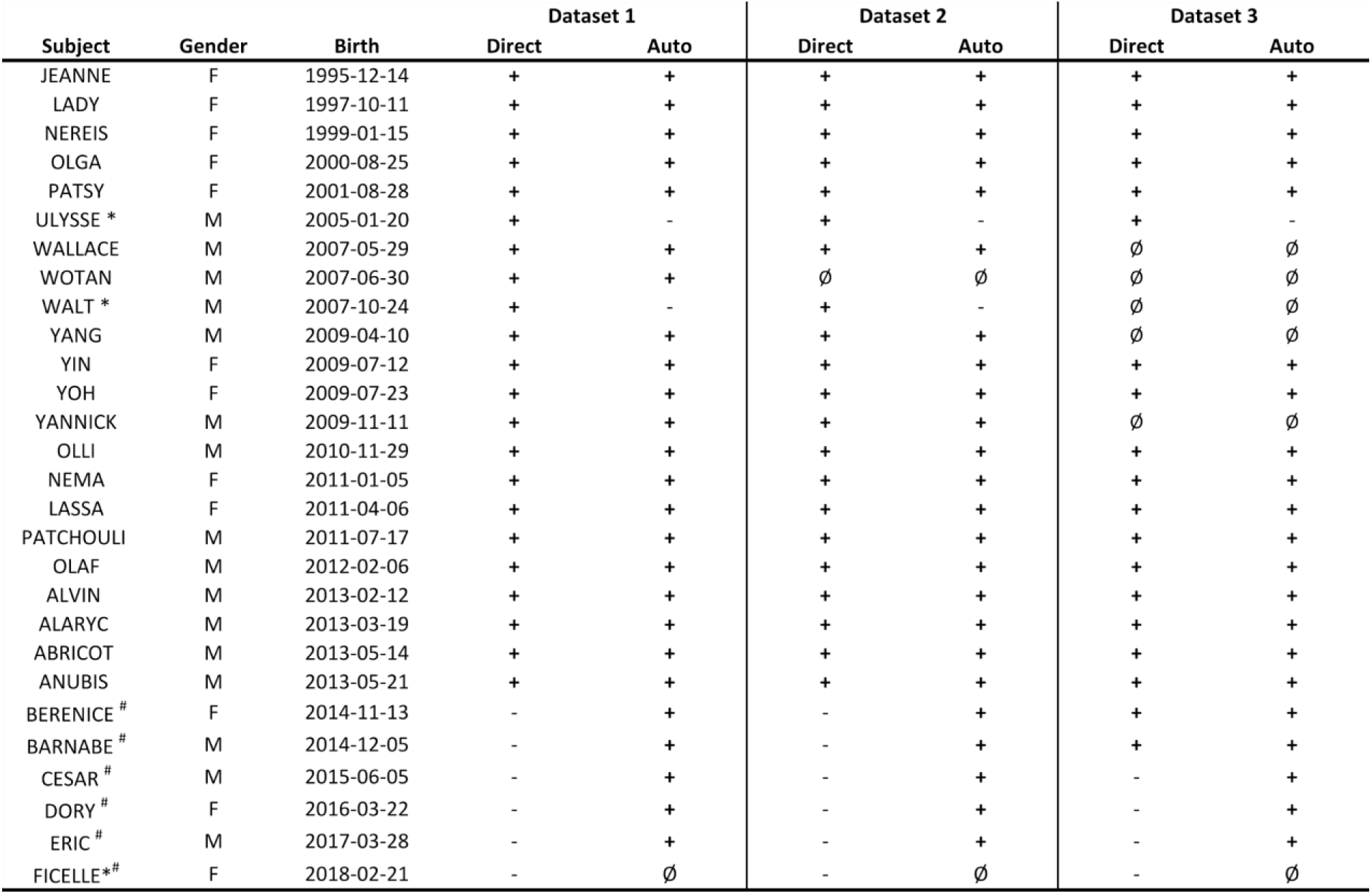
Demography of the group and subject presence (**+**), absence (Ø) or exclusion (-) in each dataset. * Corresponds to subjects that could not be included in social hierarchy measurement based on MALT conflicts. ^#^ Corresponds to subjects that could not be included in social hierarchy measurement based on direct observations of spontaneous agonistic interactions.

### Ethics

Observations were conducted non-invasively and approved by the ethical committee of the Primate Center of the University of Strasbourg, which is authorized to house non-human primates (registration n°B6732636). The research further complied with the EU Directive 2010/63/EU for animal experiments.

### Collection of direct behavioral observations by human observers

Direct behavioral observations were collected using focal animal sampling (Altmann 1974) between March 2018 and May 2019, first between the 14/03/2018 and the 29/05/2018 by one author (BS; Dataset 1) and between the 30/05/2018 and the 13/12/2018 by another author (FM; Dataset 2). Inter-Observer-Reliability was calculated during an entire week of behavioral observations (total of 89 focal follows). The outcome was Cohen’s κ = 0.89 for the recorded agonistic events and the identities of the observed individuals. Occurrences of agonistic and submissive behaviors were recorded *ad libitum*. Only data occurring in the ark and the outside shelter, where the animals were clearly in view, were recorded. Behavioral observations lasted 10 min per focal individual and were evenly spread between mornings and afternoons, from 8:30-13:00, and from 13:00-18:00. Agonistic behaviours included threats (e.g. open mouth threat), displacements (i.e. a macaque approaches another who departs immediately, e.g. at a food source, around a consorted female), chases, and physical conflict (e.g. bite, slaps). Submissive behaviours, in the context of agonistic interactions only, included facial expressions (e.g. silent-bared teeth), fleeing, crouch and screams (based on the social repertoire of Tonkean macaques described by Thierry and colleagues (Thierry et al. 2000). For each aggressive interaction, the actor and receiver were recorded, as well as if the interaction involved retaliation. In case A attacked B and B retaliated, i) with no clear winner, we encoded A-B and B-A as two independent winner-loser entries in the conflict matrix and ii) after the fight A won, we encoded A-B and B-A and A-B as three independent winner-loser entries in the conflict matrix. Behaviors were recorded using the Animal Pro Behavior software (Newton-Fischer, University of Kent 2012) on an IPod Touch (Apple), or manually on paper. The last set of direct observations was performed by another author (JW; Dataset 3) using the same focal animal sampling procedure between the 28/01/2019 and the 27/05/2019. This third dataset was already used in another study (Whitehouse and Meunier 2020).

### Automated social data using MALT

Automated data were collected at four MALTs, which the monkeys could access directly from their living environment. At the time direct observations were conducted, several cognitive tasks were available to the macaques at the MALTs, presented via a touchscreen interface. These tasks have already been described in detail (Fizet et al. 2017) and are not directly relevant for the present study. The MALTs were designed and developed at the Primate Center of the University of Strasbourg. Their development was inspired by Fagot and Paleressompoulle’s Automated Learning Device for Monkey (Fagot and Paleressompoulle 2009). These modules were set up in a shelter that was placed alongside the macaques’ enclosure. Each MALT was accessible freely 24/7, except for two-hours cleaning and refill sessions, at least twice a week. The four MALTs were placed in the same room, but were visually separated from each other by opaque Trespa^®^ boards. Monkeys were rewarded at the device for a correct answer by receiving a sip of liquid reward (2 seconds of reward, corresponding to 1 mL of diluted syrup, 1/10). MALT allows automatic identification of each subject thanks to a RFID dual-detection system (Pebayle et al. 2016). For that purpose, subjects were all equipped with two RFID microchips (UNO MICRO ID / 12, ISO Transponder 2.12 * 12mm), injected into each forearm during the macaques’ veterinary health check under appropriate anesthesia, to individually identify them when using MALT. When the RFID chip of an animal is detected, it resumes his/her personal experimental sessions, which remains open for 30 seconds after the last screen touch or RFID detection. If another animal tries to engage with the cognitive tasks while another individual’s session is active (see supplementary videos), a conflict (including which individual was replaced by who) is recorded in our database (hereafter: MALT conflict).

We considered three datasets corresponding exactly to the direct observations periods: the first dataset spanned from the 14/03/2018 to the 29/05/2018, which represents 10 257 working sessions and 995 MALT conflicts (362 remaining after filtering, see **Fig 1** and **Fig S1**); the second dataset spanned from 31/05/2018 to the 13/12/2018, which represents 62 887 working sessions and 8146 MALT conflicts (2585 after filtering); and finally, the third dataset spanned from 28/01/2019 to the 27/05/2019, which represents 30 511 sessions and 4535 MALT conflicts (1505 after filtering).

**Figure 1.**
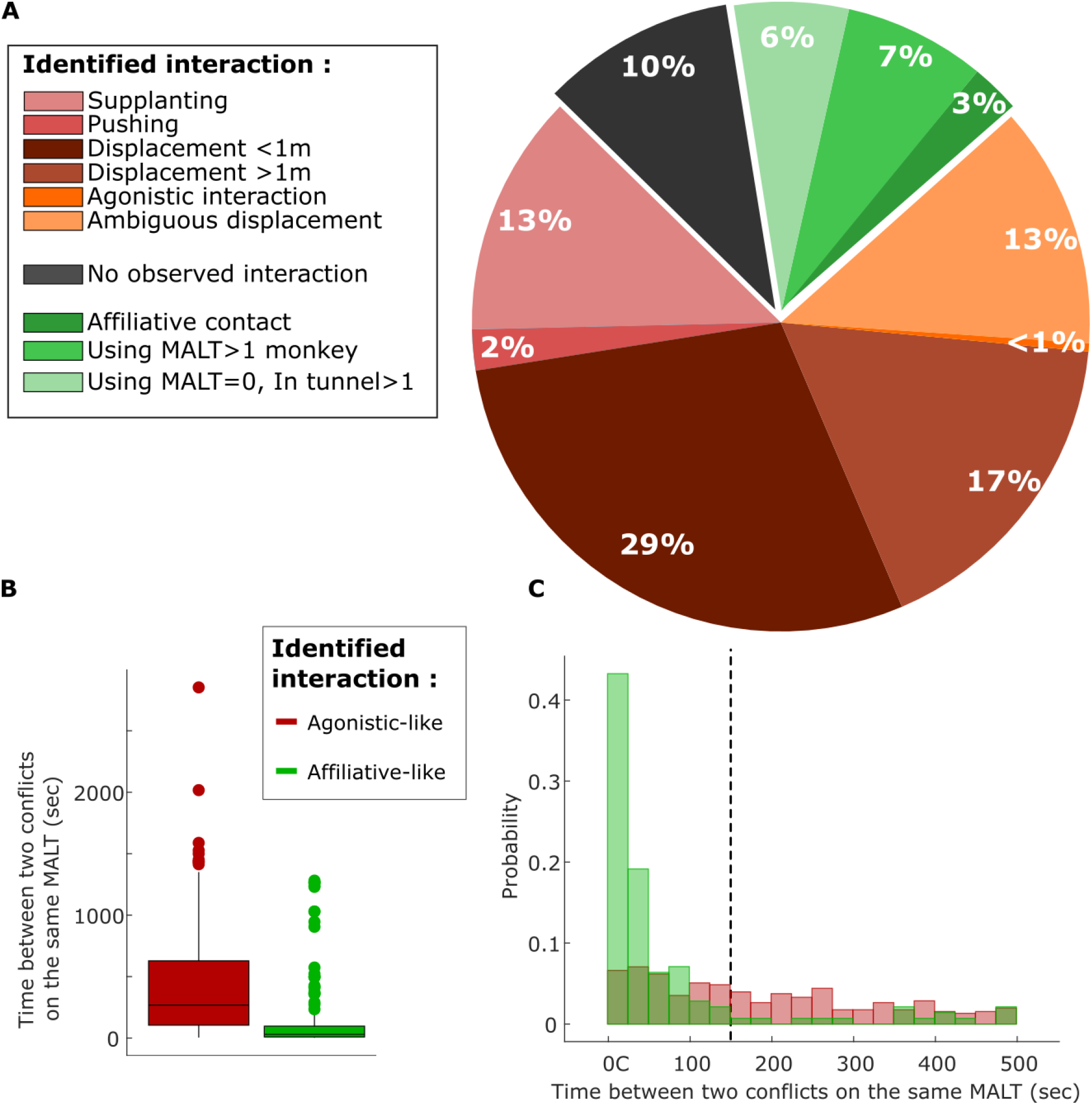
(**A**). Results of the manual scoring of the behavior of monkeys around the time of session conflicts at the MALT. The redish portions of the pie (74.5% of the sample) indicated agonistic events where a social conflict corresponding to a displacement of one monkey by another has been identified in the video recording. Dark-gray portions (10% of the sample) indicate situations where no clear social conflicts could have been identified. The greenish portion of the pie (15.5% of the sample) represent affiliative events where monkeys tolerate each other and may display affiliative behaviors. “Using MALT>I monkey” means that more than one subject was using the device; “Using MALT=0”indicates that no subject was using the device, the detection of the RFID chips of the subject happened during social interactions (usually play behavior) between animals in the tunnel linking the device to the wooded park. (**B**) The time between two conflicts on the same MALT was significantly different between affiliative and agonistic interactions (Wilcoxon rank sum, p<0.001). (**C**) Histogram representing the time between two conflicts on the same MALT for affiliative and agonistic interactions. In order to properly remove affiliative interactions from the dataset a parameter analysis helped us to select an appropriate threshold (here all conflicts that were separated by less than 150 sec were discarded). This parameters analysis for data filtering was performed by considering the mean of all correlation results for all datasets and different filtering limits (from 0 to 300 sec, see **Fig S1**). Note that the mean correlation coefficient without filtering was equal to 0.81 while the best value after filtering was equal to 0.84. Filtering affiliative and chance driven events thus marginally improved the relevancy of this measure.

### Assessing the reliability of the automated method using video scoring

Each MALT was equipped with video cameras (Microsoft LifeCam HD-3000). The video streams were cut into sections of 15 minutes each, and were automatically saved to a database if the recording contained one or more trials. We extracted and visually analyzed these video streams around the time of session conflicts. A total of 703 randomly selected videos were manually scored using The Observer® XT 10.1.548 NOLDUS software as follows. We measured four different time points for each session conflict: (1) the contester enters the tunnel area leading to the MALT, (2) the contester takes control over the MALT, (3) the former player decides to leave the MALT (body facing away from the MALT touchscreen) and (4) the former player exits the tunnel area. These time points were used to ease and control the quality of the categorization of different conflict situations (such as ‘*Displacement <1m*’ and ‘*Displacement >1m*’). Other social situations were scored based on the observed interactions between the player and the conflict monkey (e.g. ‘*Pushing*’, ‘*Supplanting*’, ‘*Affiliative contact*’). Supplantation implied the contester displaced and took the place of the former player involving physical contact but no push with hand or body part between the two monkeys. We recorded Affiliative contacts, as defined by Thierry and colleagues (Thierry et al. 2000). In order to filter events that did not represent genuine social displacement, we used the time (1) between former player departure and contester session opening and (2) between consecutive MALT conflicts (**Fig 1**). The first filter aimed at discarding chance-driven MALT conflicts (such as when one player stopped working and, shortly after, another one started using the same MALT). The second filter aimed at removing MALT conflicts that were triggered by affiliative contacts (**Fig 1**). Optimal values for these filters were empirically determined by variating these time periods and measuring the correlation between dominance hierarchy based on direct and automated methods observations (**Fig S1**).

### Data analysis

Dominance hierarchy was assessed using the David’s Scores (de Vries et al. 2006) and Elo-Rating (Neumann et al. 2011) both using the package ‘EloRating’ in R (R Core Team 2014). Both the use of David’s Score and Elo-rating for the assessment of hierarchical structure is common throughout the study of animal behaviour (Neumann et al. 2011) and therefore we chose to consider both methods here. One of the main differences between these two approaches is that David’s score is calculated on a complete interaction matrix, where the temporality of interactions is not considered, whereas Elo-rating is calculated based on sequence of events where the order of interactions is important and taken into account. This provides Elo-rating with the added benefits of being able to assess the dynamics of a hierarchy across time, and allowing for the extraction of hierarchy data at specific time points. In all cases where Elo-rating was used, we first optimised the k factor (i.e. the maximal amount of ‘points’ an individual can get from an interaction, function: optimizek, package: Elorating). We assessed the correlation between dominance hierarchy based on direct observations and our automated method using Spearman’s and Pearson correlation test for each dataset separately. Non-parametric correlation method was indeed used when considering ordinal ranks. Sample sizes were 20, 19, and 18 individuals for dataset 1, 2 and 3, respectively. Analyses were performed using custom scripts in Matlab (R2018a, the Mathworks), R scripts were called using Matlab (Weirong Chen 2020) and Gramm graphical toolbox was used (Morel 2018).

### Application of automated data: a proof of concept

As an example of how such automated data could be applied, we firstly calculated the Elo-rating of our group across all of our automated observation periods so far, totalling 1095 consecutive days (02/02/2017 to 31/03/2020). During this period, the MALT recorded 23878 conflicts. Two key events could represent a disruption to the hierarchy - the removal of one mid-ranking adult male in the group (event 1, 26/05/2018), and the removal of four high-ranking adult males in the group (event 2, 18/01/2019). In this species, adult males often migrate to neighbouring groups. Here, the decisions to remove these animals were in order to mimic this natural change in macaques’ group dynamic (Riley 2010) and to ultimately avoid the potential for inbreeding. To assess the effect of these removal events on the hierarchy, we extracted day-by-day stability of the hierarchy (function: stab_elo, package: Elorating), which provides us with a score between 0 and 1 for each day (where 1 represents a stable hierarchy with minimal rank reversals). Using this data, we compared the stability of the hierarchy in the 50 days prior to an event, and compared that with the stability of the hierarchy in the 50 days after an event. These pre-and post-event data points were then compared with a Wilcoxon signed-rank test.

## Results

The dominance hierarchy of a group of semi-free ranging Tonkean macaques was assessed using two independent approaches. Data collected with classical direct observations of the animals’ agonistic interactions (n=948), was compared to an automated method, based on social displacements when using MALT (n=13 676) during the same period of time.

We analyzed 703 randomly selected videos recorded around the time of the MALT conflicts. We scored the type of interactions between the subjects (See Methods and Supplementary Videos) and found that in at least 74.5% of the cases, the MALT conflicts included an agonistic interaction (**Fig 1a**). These social interactions included different active forms of social displacements such as supplanting or even pushing the former user of the MALT. We also observed ‘Ambiguous displacement’ when more than two individuals were involved in the conflict, or the conflict had no clear outcome (e.g. the displaced individual did not leave the area). In the 10% of the cases, no social interactions were detected at all, as the player left the area and, within the next 30 seconds, another individual came to use the MALT. These situations were arguably driven by chance even if we cannot exclude that auditory or visual cues, which here cannot be detected by the human observer, prompted the animal to leave the MALT (see ‘no observed interaction’ in **Fig 1a**). In 15.5% of the cases, MALT conflicts were related to affiliative interactions. These include situations such as young subjects, playing around the MALT, accidentally detected within the same 30 seconds windows, which created a conflict on the MALT (see ‘using MALT =0, tunnel>1’ in **Fig 1a**). We also recorded co-presence inside the tunnel without any sign of agonistic interactions (see ‘using MALT >1’ in **Fig 1a**). For instance, one individual was observed working while the other was drinking the juice reward. Such situations can be regarded as interesting co-working and/or co-feeding tolerance examples and may require further investigation (Carne et al. 2011; Dubuc et al. 2012).

We aimed to correct the 25.5% of estimated recordings that did not represent any form of displacement (based on the video analysis above). To discard chance-driven MALT conflicts, we considered the delay between the departure of the first player from the MALT (last touch recorded on the touchscreen) and the MALT conflict (reading of the RFID chip of the next monkey). A threshold of 20 seconds gave the best correlation coefficient between rankings based on automated and direct observation (**Fig S1, Fig 2**). MALT conflicts triggered by affiliative interactions, such as individuals playing around the MALT, are not directly relevant for assessing dominance hierarchy in NHP and should thus be excluded from the analysis. With several individuals present in the tunnel or next to a MALT, the frequency of MALT conflicts is likely to increase. We thus measured the time between two conflicts on the same MALT and showed that it was significantly different between agonistic and affiliative interactions (**Fig 1b**, medians respectively equal to 282 seconds and 30 seconds, Wilcoxon rank sum, p<0.001). We empirically found the best threshold in order to filter our data (**Fig S1** and **Fig 1c**). MALT conflicts were discarded if they were separated by less than 150 seconds.

**Figure 2:**
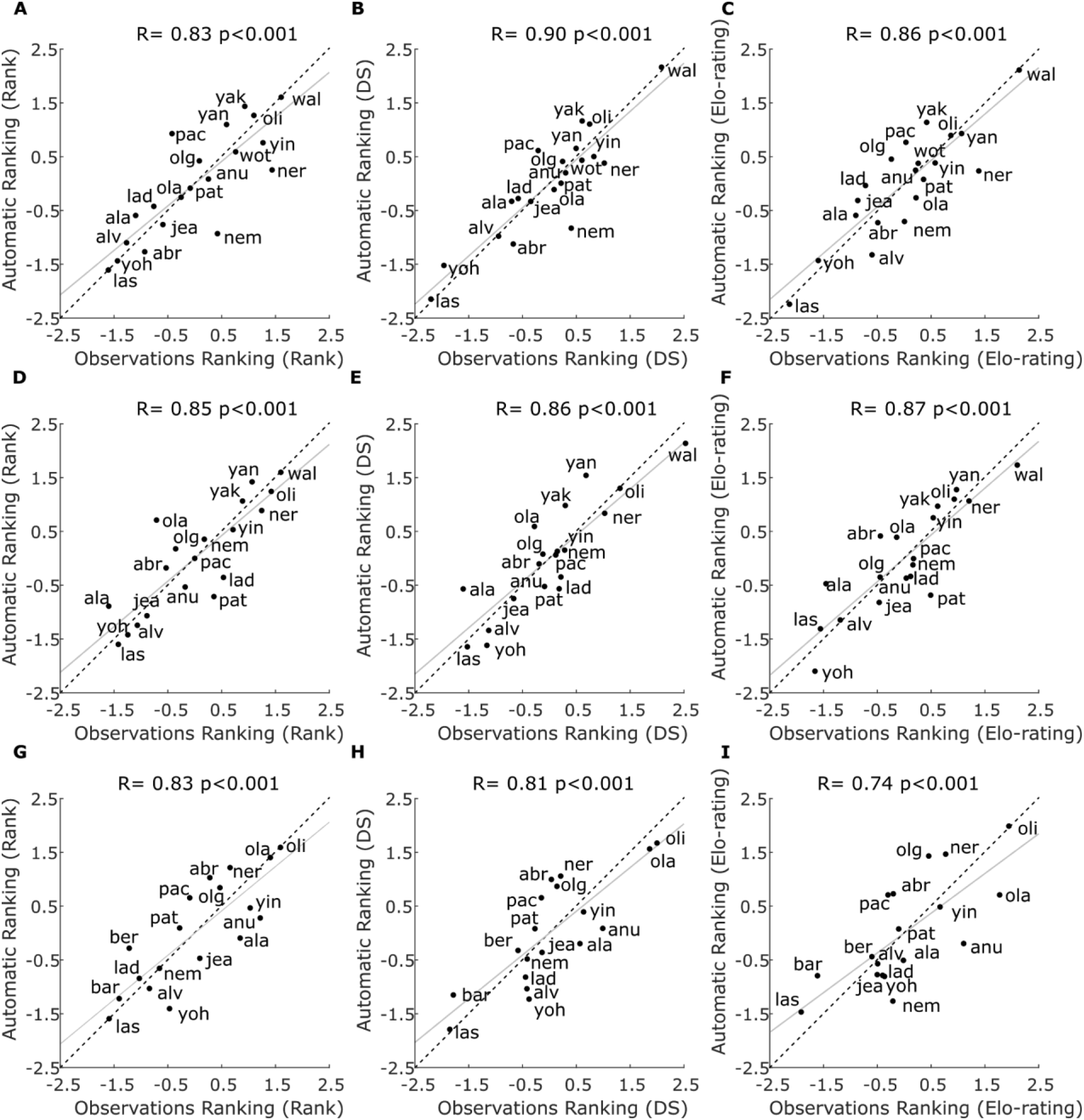
Comparison of social hierarchies computed using direct observations of behaviors (Direct Observations Ranking) and MALT conflicts (Automatic Ranking) in three datasets (rows) and using three different measures of social hierarchies (columns). For all panels, the gray line represents simple least squares regression and the dashed line the reference. Each row represents a given dataset analysis and each column a different method to compute the social hierarchy. In panels (**A,D,G**), the social hierarchies were calculated using the ordinal ranks obtained with the David’s Score (DS); Correlation coefficient R and p values corresponded to Spearman rank correlations. In panels (**B,E,H**), DS values were used; Correlation coefficient R and p values were from Pearson correlation. In panels (**C,F,I**), Elo-ratings were considered; Correlation coefficient R and p values were from Pearson correlation. Sample sizes were 20, 19, and 18 individuals for the dataset 1(**A,B,C**), 2(**D,E,F**) and 3(**G,H,I**), respectively. For graphical purposes only, all data were z-scored.

Using the two above-mentioned filtering parameters 67.5% of all recorded MALT conflicts were removed from the dataset. We found substantial correlations between the David’s Scores (DS ordinal rank and values, **Fig 2adg** and **Fig 2beh**, respectively) and Elo-rating (**Fig 2cfi**) computed with direct observations and MALT conflicts (**Fig 2**, Spearman rank and Pearson correlation all R>=0.74 and p<0.001; overall mean R = 0.84). Note that unfiltered data gave very similar results (overall mean R = 0.81 and all p <0.01). In every case, the hierarchy was linear (Permutation test, p<0.001 (de Vries 1995)). Overall, these analyses demonstrate that conflicts occurring during the use of the MALT represent a good proxy of social conflicts.

Finally, we used more than 3 years of MALT conflicts to compute the hierarchy of the group (**Fig 3a**) and we considered the impact of male-removal events on the stability of the hierarchy (**Fig 3ab**).These analyses showed a significant reduction in stability after the removal of a mid-ranking male (**Fig 3c**, W = 837, p = 0.003), but no significant changes after the removal of four highest-ranking males (W = 1342, p = 0.7831).

**Figure 3:**
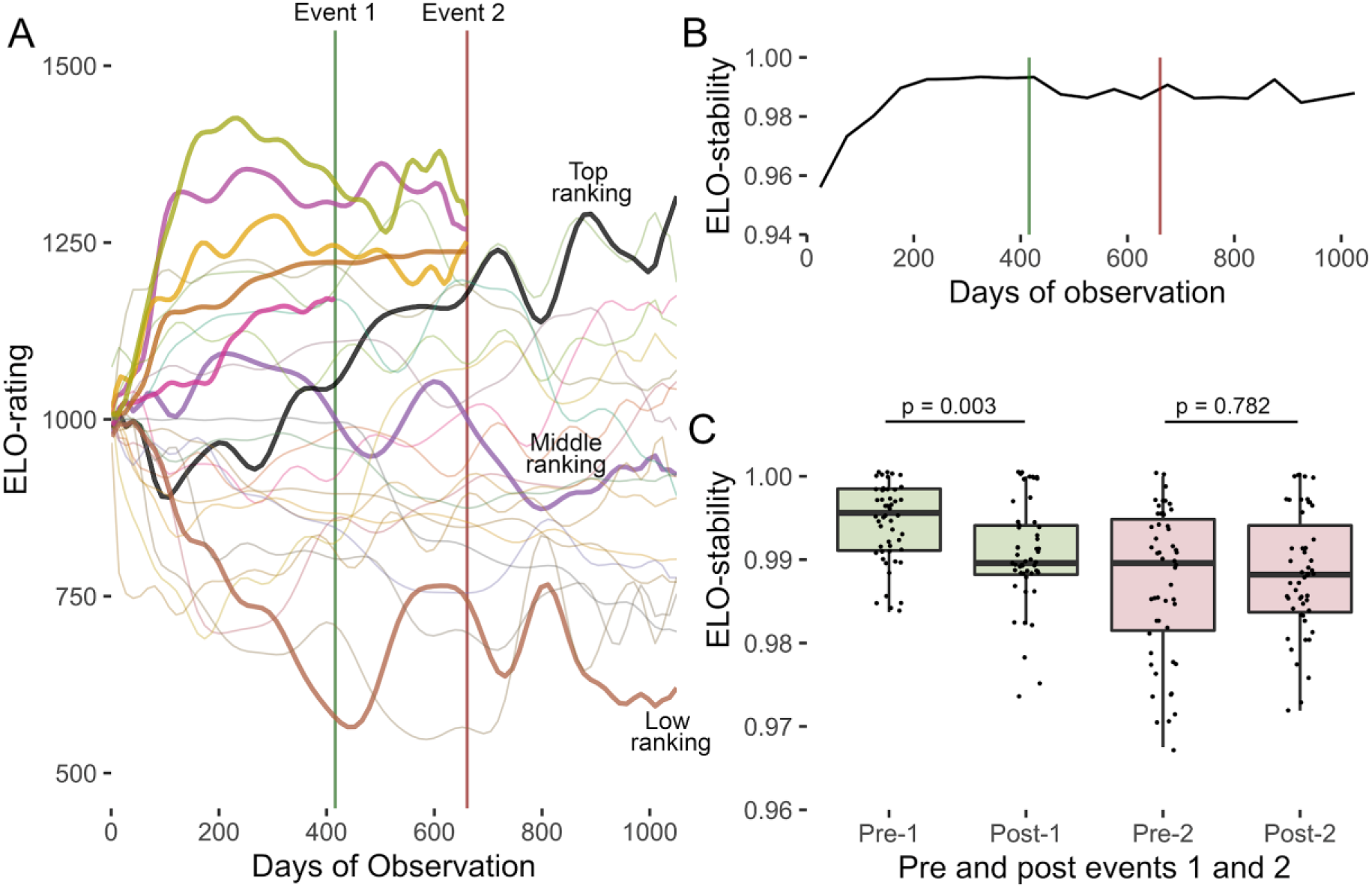
Proof of concept: automated Elo-rating across all observation periods and the effect of animal-removal events on the hierarchy. (**A**) Elo-plot across time; smoothing has been applied to each line for visibility and a number of key animals have been highlighted with bold lines. All of the males that were removed during the 2 events (the timing of events are visualised with vertical lines) are highlighted here (n=5), in addition to the animals that were calculated as the highest, lowest, and most mid-ranking at the end of the observation period. (**B**) The stability of the hierarchy across time. Here the stability data is presented in average blocks of 50 days for visualisation purposes. (**C**) Boxplot visualising the data used in our analysis (1 data point = 50 days pre- and post-male removal events. Data for event 1 can be seen in green, data for event 2 can be seen in red.

## Discussion

Social hierarchies can be measured based on the outputs of dyadic conflicts over access to any resources (Hamilton 1960; Boelkins 1967; Christopher 1972; Clark and Dillon 1973; Chamove 1983). Only few solutions already exist in order to measure dominance interactions in animals automatically (Hrolenok et al. 2018; Evans et al. 2018). In this study, we considered several months of daily use of MALT by 25 semi-free ranging Tonkean macaques in order to assess the dominance hierarchy of this group. Our method does not require human observers and in theory, only one MALT is needed to achieve such a measure of the hierarchy within a social group. In a comparable amount of days, MALT can record about 10 times more conflict events compared with direct sampling methods by human observers. While our analysis between direct and automatic data reveals a strong agreement in the hierarchical structure, some difference remains (Evans et al. 2018; Hrolenok et al. 2018). Several points can be considered to explain these differences. First, ethological sampling cannot assume to be completely error-free. For instance, inter-rater reliability analysis achieving 80% congruence is usually considered as acceptably high-agreement (McHugh 2012). Note that we found an overall mean correlation coefficient between automatic and ethological ranking of R=0.84, which is about what would be expected when correlating the same two measurements that each contain 20% of independent noise. That being said, our automated method also has limits and some of the discrepancies between observation and automatic measurements may be due to different social contexts where conflict arises (Brennan and Anderson 1988). For instance, MALTs are preceded by a tunnel (of approximately one meter) that promotes face-to-face interaction that may impede coalition formations. In addition, the motivation of individuals to use the MALT (that integrates, at least, the value for diluted syrup rewards and the subjective cost of performing cognitive tasks) may also come into play in the decision of an animal to compete or not with another. These variables may not influence other types of social conflicts that are used to measure social hierarchy during direct observations. Human observers can record a number of context elements that may be especially useful for some research questions (e.g. formation of rank leveling coalitions). The use of MALT to assess dominance hierarchies is therefore limited by the lack of fine grained information about the context under which naturally occurring conflicts arise (e.g. for access to fertile females) and the possibility for bystanders to intervene (Petit and Thierry 1994). In addition, if they are not interacting enough with the MALT, some subjects cannot be included in this measurement of the dominance hierarchy (here n= 2 / 28 subjects). On the other hand, this automatic method allowed us to record information that was difficult to obtain using direct observations. In particular, we were able to gather dominance data from five juveniles that were considered during direct observations due to challenging subject identification. MALT can thus also be used to assess the hierarchy between juveniles which is often neglected in other studies (Fedurek and Lehmann 2017). MALT may thus also provide new information on the role of juveniles in a species social organization, or allow for a detailed assessment of the development of the social rank of juveniles over time.

It should be noted that no ethogram is required to use this method and as long as displacement is considered agonistic, it could be theoretically used in any animal. Hence, even if we demonstrate the relevancy of this method in a single species of NHP, it seems parsimonious to think that this can be safely generalized to other NHP species. In this tolerant species of macaque, we estimated that about 15.5% of the conflicts detected with the MALT might not represent social conflict events. For instance, manual scoring of a subset of MALT conflict videos revealed unexpected situations when macaques appeared to “share” a device (See Supplementary Videos), i.e. one individual collecting the reward of the other one. Tonkean macaques are known to be more socially tolerant than other species of monkeys (Thierry 2007) and these affiliative events are thus likely to be more rare in other species of NHP. In any case, thanks to an appropriate filtering criterion, we achieved to remove most of MALT conflicts that might not represent an agonistic interaction. However, we noted that the presence of these affiliative events in the dataset was only marginally impeding the measure of social hierarchy (differences of 0.03 between the mean correlation coefficients of unfiltered and optimally filtered data). The close affiliative interactions observed in the MALT are likely restricted to a few preferred social partners. Being able to also identify individuals that are around, but are not directly using the MALT, could reveal a social tolerance or affiliation network that can be related to dynamic coalitions formation (Berghänel et al. 2011). Further development of this method may include face recognition to achieve such goal (Krause et al. 2013; Witham 2018; Zhang et al. 2018; Schofield et al. 2019). Generally, further development is needed to reliably and automatically assess the many dimensions of the affiliative networks of NHPs, but this is beyond the scope of this present study.

One of the strengths of this automated measure is the continuous recording events generating a much bigger dataset than human observations. For instance, as a proof of concept, we used this automated approach to assess the effect of two events of male-removal on the hierarchical stability of the group. This analysis considered more than 1000 days of observations, this is, to the best of our knowledge, not the longest (Rhine et al. 1989; Rhine 1994; Goldman and Loy 1997; Robbins et al. 2005) but the most detailed assessment of social hierarchy in a group of NHP that has ever been reported. Interestingly, removing a mid-ranking male caused an immediate reduction in group stability, however, removing four high-ranked males had no significant impact. This shows that the number of individuals removed (or migrating) from a group can be less influential than the positions they hold in the social network. Indeed, middle-ranking males represent key nodes in the organisation of the dominance hierarchy, as they could form coalitions with either the alpha to reaffirm its dominance, or participate in rank reversal coalitions against higher-ranking males (van Schaik et al. 2004). Consistent with our observations in Tonkean macaques, patterns of grooming associations in captive crested macaques remained unchanged after the removal of seven individuals, mainly adult males, whereas the introduction of a single new adult male triggered an increase in grooming activity among females (Cowl et al. 2020). However, these observations are based on two single cases and should be treated with caution. More importantly, these data provide an example of the potential applications of continuous and automated conflict data that could ease captive group management and pave the way for a better understanding of NHP social dynamics.

Overall, we report that the presence of food rewards (here flavored syrup diluted in water) accessible through the correct usage of MALT creates a competition over this resource which induces dominance behaviours in the macaques. We show that the social hierarchy computed thanks to these social displacements was highly consistent with the one computed using observation of spontaneous social conflicts in the monkeys’ living environment. Our analysis further suggests that the presence of affiliative events is not dramatically impeding the relevancy of these automatic measurements, likely thanks to the considerable volume of genuine social displacements that can be recorded by this method. Our study clearly supports the use of MALT to automatically, reliably and longitudinally assess the dominance hierarchy of NHPs.

## Acknowledgements

The authors are grateful to the University of Strasbourg and Silabe (https://silabe.com/) for supporting this research and providing expert animal care. We also would like to thank Adam Rimele for computer architecture and programming support. The MALT development was supported by the University of Strasbourg Institute for Advanced Study (USIAS) as part of a USIAS fellowship to HM.

## Supplementary Material

Supplementary videos are available at: https://seafile.unistra.fr/f/734a7bae4ff44c96b5b4/

---

**Fig S1.**
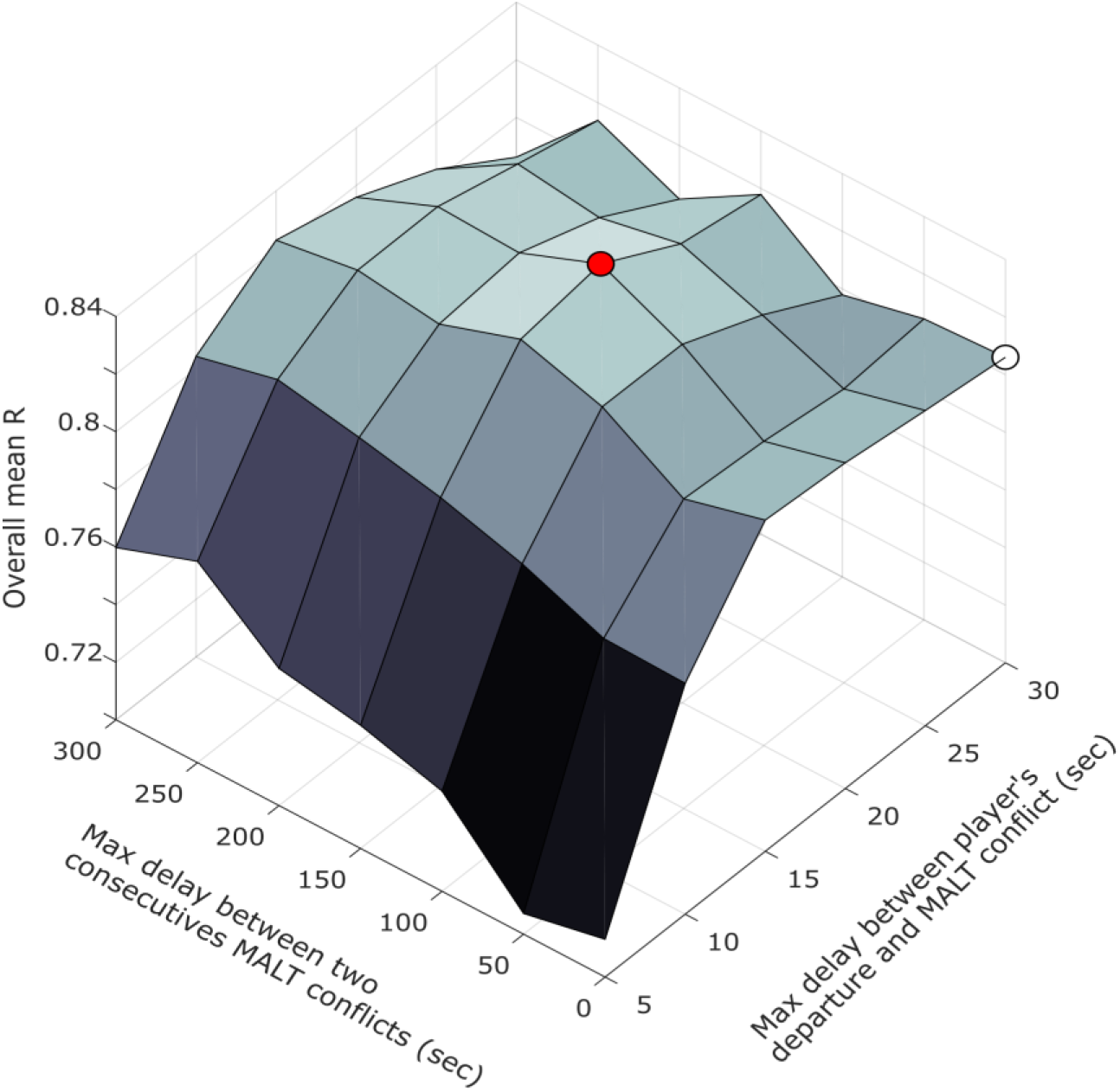
Optimisation procedure for the two parameters used to filter MALT conflicts. Analysis of the videos of MALT conflicts revealed the presence of chance driven events (absence of social interaction) and events containing affiliative interactions between the player and the conflict monkey. These events did not represent a genuine social displacement and should thus be logically discarded to compute hierarchy of dominance. Absence of social interaction can be filtered using the delay between the departure of the player from the MALT (last touch recorded on the touchscreen) and the MALT conflict (reading of the RFID chip of the conflict monkey). Affiliative events can be filtered based on the delay between two consecutives conflicts on the same MALT. Overall mean R value is the mean of all correlation coefficients for all datasets. The white dot corresponds to unfiltered data (30 sec and 0 sec, R=0.84); the red dot corresponds to the value of the filtering parameters that gave the best correlation coefficients between automatic and direct observations (here 20 sec and 150 sec, R=0.81).

